# Getting More by Asking for Less: Linking Species Interactions to Species Co-Distributions in Metacommunities

**DOI:** 10.1101/2023.06.04.543606

**Authors:** Matthieu Barbier, Guy Bunin, Mathew A. Leibold

## Abstract

One of the more difficult challenges in community ecology is inferring species interactions on the basis of patterns in the spatial distribution of organisms. At its core, the problem is that distributional patterns reflect the ‘realized niche’, the net result of a complex interplay of processes involving dispersal, environmental, and interaction effects. Disentangling these effects can be difficult on at least two distinct levels. From a statistical point of view, splitting a population’s variation into contributions from its interaction partners, abiotic environment and spatial proximity requires ‘natural experiments’ where all three factors somehow vary independently from each other. On a more conceptual level, it is not even clear how to meaningfully separate these processes: for instance, species interactions could depend on the state of the abiotic and biotic environment, and these two processes may combine in highly non-additive ways. Here we show that the latter issue arises almost inescapably, even in a simple theoretical setting designed to minimize it. Using a model of competitive metacommunity dynamics where direct species interactions are assumed to be context-independent, we show that inferring these interactions accurately from cross-species correlations is a major challenge under all but the most restrictive assumptions. However, we also find that it is possible to estimate the statistical moments (mean value and variance) of the species interactions distribution much more robustly, even if the precise values cannot be inferred. Consequently, we argue that study of multi-species spatial patterns can still be informative for theoretical approaches that build on statistical distributions of species parameters to predict macroscopic outcomes of community assembly.

---

A central issue in community ecology is to identify which processes and mechanisms are most important in determining species presence and abundance across space and time in given communities. While this can be accomplished by carefully designed experimental methods, this is often logistically impractical and indirect inference is used: we attempt to infer parameter values for a **process**-based model via the analysis of naturally-occurring **patterns** of species distributions. While early work focused on the direct analysis of co-distributions (i.e. correlations) in the abundances or occurrences of species, there has, over time, become apparent that this can be misleading and an increasingly sophisticated set of analytical tools that try to do this has emerged.

To give context to this issue, we examine the challenges in using distribution patterns to evaluate species interactions in the critiques and subsequent debate in response to Diamond (1975)’s ‘Assembly Rules’ that were proposed to explain coexistence patterns in relation to interspecific competition in community and biogeographic data. Diamond (1975)’s assembly rules focused especially on looking at the significance of negative co-distributions in community patterns among potentially competing species (i.e. so called ‘checkerboard’ patterns of coexistence). While a long-lasting debate ensued on the statistical significance of these patterns (Connor & Simberloff, 1979; Gotelli & McCabe, 2002) and the use of ‘null models’ (Gotelli & Graves, 1996), few, if any, questioned the basic hypothesis that negative co-distributions were in fact robust indicators of competition until much later.

More recent work has increasingly recognized the confounding roles of environment, disturbance, isolation and dispersal and proposed more sophisticated methods for the study of co-distributions, e.g. (Patterson & Atmar, 1986; Leibold & Mikkelson, 2002; Peres-Neto *et al*., 2006). These issues have been resurfacing especially in microbial ecology due to a tide of data (Berry & Widder, 2014). Additionally, there has developed a broader focus on other types of species interactions (e.g. Cazelles et al. 2015). The analysis of species interactions in general, and accounting for environmental, space and dispersal have been merged by the more recent development of more sophisticated methods such as joint species distribution models e.g. (Ovaskainen *et al*., 2017) and related methods that aim to partition distribution patterns in relation to ‘abiotic’ (environmental), ‘biotic’ (involving interactions among species) and ‘movement’ (involving spatial effects mediated by dispersal effects). While the sophistication of such methods is rapidly developing, especially in addressing computational and statistical issues, this work is increasingly revealing that inferring process from pattern is not trivial (Barner *et al*., 2018; Thurman *et al*., 2019; Blanchet *et al*., 2020; Poggiato *et al*., 2021).

In parallel with these developments in biogeographic ecology, issues have arisen in work that is more narrowly focused on inferring species interactions from relative abundance patterns in a homogeneous setting (Levine, 1976; Lawlor, 1979). That work ignores confounding effects of environment and dispersal, usually by focusing on situations where these are constrained (experimentally) or assumed to be negligible (by the selection of data). It typically attempts to infer pairwise interactions by comparing abundances across different overlapping sets of species, see e.g. (Barbier *et al*., 2021). Two questions have been at the heart of that work: what do we mean by species interactions, and do they depend on their biotic context? We wish to argue here that these two questions are connected, and that ecological dynamics inescapably give rise to some context dependence in species interactions.

To clarify, interactions in the statistical sense can be readily observed in species abundance patterns, but from a causal point of view, abundances do not directly affect each other. Rather, abundances control processes (e.g. inducing mortality in a competitor), and these processes in turn determine changes in abundances in the short or long term. The former have been termed ‘direct effects’, and the latter ‘net effects’ (Lawlor, 1979). Even if the direct effects of one species on another’s dynamics were context-independent, its net effects on abundances in the long term can be mediated many other species, along chains of indirect impacts playing out over time, and can thus depend on the whole community’s composition (Schaffer, 1981; Zelnik *et al*., 2024). We must therefore carefully define direct and net effects and specify which we are trying to infer. It has been proposed that co-distribution patterns across space can be used to deduce a fixed matrix of net effects between species (Ovaskainen *et al*., 2017), but this assumes that species composition does not change over the metacommunity. This method is consequently not likely to be sufficiently robust to apply except under very highly controlled or limited conditions.

Nevertheless, the analysis of distribution and co-distribution patterns of species in metacommunities suggests that, while obtaining exhaustive parameter estimates is still challenging (e.g. (Blanchet *et al*., 2020; Poggiato *et al*., 2021; Leibold *et al*., 2022), some broader features can be related to the processes that generate them (Ovaskainen *et al*., 2017; Ovaskainen & Abrego, 2020; Leibold *et al*., 2022). For instance, it may be possible to obtain good estimates of the *average* competition strength in a community (Fort, 2018). A body of theoretical work on so-called “disordered systems” (Barbier *et al*., 2018) proposes that, when biotic interactions are numerous enough and sufficiently independent from each other (unstructured, contrary to e.g. a competitive hierarchy), they can be treated as random-like, and only their mean and variance matter in determining outcomes of community assembly such as diversity or stability. Inspired by this theory, we hypothesized that statistical features (e.g. mean and variance of either direct or net effects) may be a more robust inference target than the precise network of interactions. We further wondered if these statistical properties might also be robust to some of the other concerns outlined above such as the existence of composition change, the amount of dispersal and degree of environmental variation.

Here, we propose to explore limits to inference that stem from the very nature of the dynamical processes, which entangle contributions from the environment and from multiple species in ways that might or might not be possible to disentangle at all. We use process-based simulations of community assembly in disordered communities (as defined above) to generate simulated data under different conditions involving species niches, environmental gradients, dispersal and interaction strength. We consider predominantly competitive interactions, because our simple Lotka-Volterra model can be unstable under strong facilitation, although we believe that many of our results would otherwise extend to any interaction type. We then infer parameters from the analysis of resulting patterns to evaluate how well they can be used under these various scenarios.

Given that previous studies have already established that biotic interactions can strongly bias our estimates of species’ environmental preferences (Poggiato *et al*., 2021), we go one step further to show that, even in a setting where we could assume a good estimate of these environmental preferences, we may still be unable to correctly extract species’ interactions from their spatial co-occurrence. Yet we find that, even when species interactions cannot be estimated in detail, it remains possible to correctly infer their community-level statistics, i.e. how strong and diverse biotic interactions are overall in the ecological dynamics of the community. This suggests that we may extract more robust information out of spatial patterns by asking for a less detailed description of the underlying processes.

## 1 Methods

To evaluate how we might infer processes involving species interaction coefficients from patterns in species distributions in a landscape or metacommunity, we considered a highly simplified modeling framework as a limiting case. We assumed that species interacted in a spatially continuous lattice in which local (within cells) interactions could be described by simple Lotka-Volterra equations connected by dispersal from nearby cells. Previous work on inferring species interactions from possible distribution patterns suggested that this could work under similar assumptions, at least under some limiting conditions) (Levine, 1976; Lawlor, 1979). We assume that more complex assumptions (e.g. non-linear interactions, heterogeneous dispersal etc), would make the inferences we are interested in even less likely and our approach should thus be seen as an ‘optimistic’ evaluation. In essence, we are asking “how well can we **hope** to do in making such inferences with current approaches?”

We intuit that a significant obstacle to inferring the details of species interactions is the covariation between the abundances of all interaction partners, and between each of them and the environmental factors: we cannot discriminate how strongly each of these factors impacts the presence of any given species if our observations do not provide ‘natural experiments’ where they vary independently (or actual experiments that impose various species compositions in the same environmental conditions (Barbier *et al*., 2021)). Therefore, we start by focusing on two highly contrasting cases. In one case, we model a scenario that is most likely to be successful in inferring interactions parameters from distributions, because many species compositions are realized for each environmental condition in the absence of local dispersal. We then contrast this with a scenario that includes dispersal that allows environmental tracking by species in the metacommunity.

Within these two scenarios, we analyze a number of simulations that vary in the mean and variance of species’ interaction strengths. We know from prior literature, e.g. Roy *et al*. (2020); Hu *et al*. (2021), that our simulation model may exhibit complex dynamics, such as chaotic fluctuations, when species interactions are sufficiently strong. To situate ourselves in the most favorable setting for inference, we avoid strong interactions or strong dispersal, so as to select a dynamical regime where species reach a globally stable equilibrium that reflects how favored they are in their local abiotic and biotic environment. In each simulation, we then attempt to infer interactions in detail, as well as derive their mean and variances.

### 1.1 Model setting

We consider a metacommunity on a 2D landscape of 64 *×* 64 pixels, each represented by a coordinate vector *x*. At each point, we model a single environmental factor *E*(*x*) whose values range in [–50, 150] (Fig. 1). We then simulate Lotka-Volterra dynamics with dispersal from neighboring patches *y*

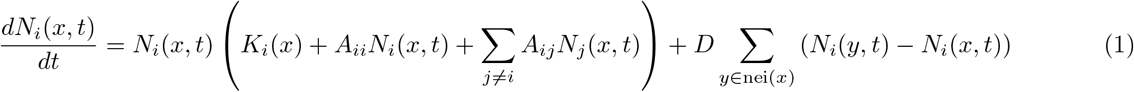

**Figure 1:**
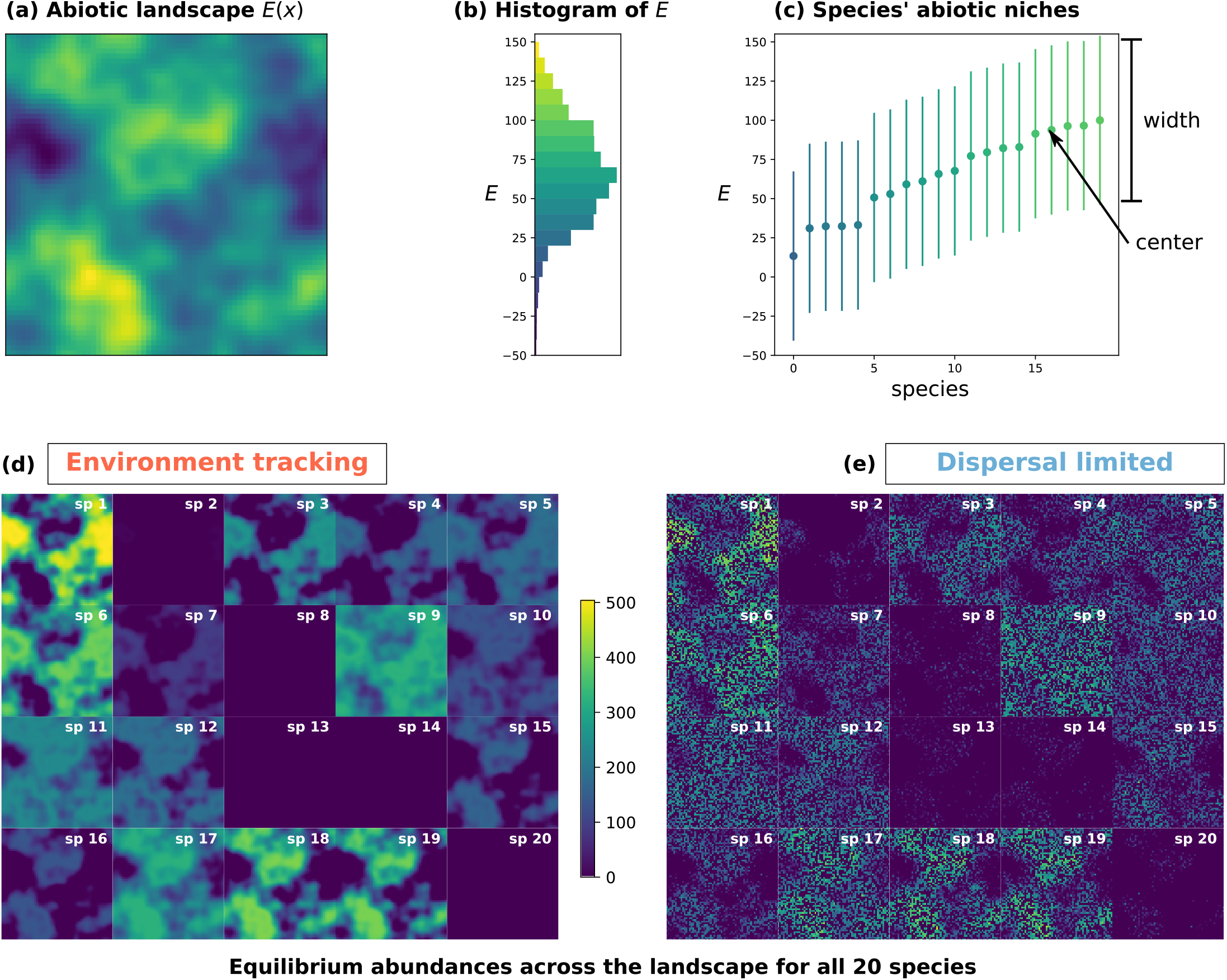
Environmental factor and final species abundances across the landscape in an example simulation run. **(a)** Landscape *E*(*x*) with *x* a two-dimensional coordinate vector (64 *×* 64 pixels called patches). **(b)** Histogram *P* (*E*) of values of the environmental factor. **(c)** Centers *c*_*i*_ (dots) and width *w*_*i*_ (bars) of abiotic niches for all 20 species. **(d-e)** Equilibrium abundances *N*_*i*_(*x*) for all species (rescaled by extinction threshold *N*_*c*_ = 10^−3^ so that values less than 1 indicate local extinction). The two colonization scenarios described in Sec. 1.2 are represented here: **(d)** environment tracking, where every species is initially seeded in every patch (or are allowed moderate dispersal), and **(e)** dispersal limitation, where species are seeded independently in 50% of patches at random, and cannot disperse. In the latter case, species that would outcompete others in a given environmental condition might be absent by chance in some patches with that condition. Thus, several species go fully extinct at the landscape level due to competition in the environment tracking scenario **(d)**, while all species are present in at least part of the landscape in **(e)**. Parameters: ⟨*w*_*i*_⟩ = 50, ⟨*A*_*ij*_⟩ = –0.3, std(*A*_*ij*_) = 0.09, *D* = 0.

Furthermore, species are considered extinct when *N*_*i*_ *< N*_*c*_ = 10^−3^, but are allowed to invade again: their growth rate *dN*_*i*_*/dt* is replaced by max(0, *dN*_*i*_*/dt*), which may allow them to regrow above the extinction threshold if this rate remains positive. At the end of the simulation, the abundances of species under the threshold are set to zero, to be ignored in our analyses.

Intra-specific competition is set to *A*_*ii*_ = –1 for all species, while inter-specific interaction coefficients *A*_*ij*_ are independent of the environment and drawn randomly for each species pair (*i, j*) from a normal distribution with prescribed mean ⟨*A*⟩, standard deviation std*A*, and symmetry sym*A* = corr(*A*_*ij*_, *A*_*ji*_). We typically take a large negative mean, so that coefficients are predominantly competitive, to avoid the breakdown of the Lotka-Volterra model with strong facilitation.

The coefficients *K*_*i*_ determine each species’ carrying capacities (equilibrium abundance in monoculture) since *A*_*ii*_ = –1, and thus *N*_*i*_ = *K*_*i*_ at equilibrium in the absence of other species and of dispersal. These carrying capacities are modelled using a unimodal “niche” function of the environmental factor:

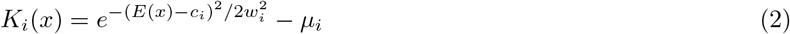

with “mortalities” *μ*_*i*_ drawn uniformly between 0 and 0.5, niche centers drawn uniformly *c*_*i*_ *∈* [0, 100] and widths *w*_*i*_ normal i.i.d. (see parameters in SI). The addition of *μ*_*i*_ ensures that *K*_*i*_ *<* 0 when the environment deviates enough from species’ optimum, i.e. species may not grow at all in sites that are too unfavorable.

As noted above, drawing on previous theoretical and numerical work (Bunin, 2017; Zelnik *et al*., 2021) we choose parameters in this model that select a global equilibrium regime: species abundances reach a stable equilibrium in each site based only on the pool of species that can access that site, their local environment and interactions. This requires that std*A* is small enough to avoid loss of stability leading to complex nonequilibrium dynamics (Bunin, 2017), and *D* is small enough to avoid significant source-sink dynamics and mass effects, i.e. situations where local abundance are strongly driven by fluxes from neighboring sites (Leibold *et al*., 2004; Zelnik *et al*., 2019). These other situations may also be of ecological relevance, but the regime considered here was the most appropriate considering the questions we wish to tackle, as argued in Discussion.

### 1.2 Colonization scenarios and dispersal

We expect that interactions can be inferred more successfully in biodiversity experiments where different species compositions are imposed in the same environmental conditions (Barbier *et al*., 2021). This suggests testing two distinct scenarios for how species are distributed in the landscape (Fig. 1):

(DL) **“Dispersal limited”** scenario where each species is only seeded (i.e. given positive initial abundance) in half of the patches at random, and cannot disperse between patches.

(ET) **“Environment tracking”** scenario where all species are initially seeded in every cell (or can freely disperse, usually setting *D* = 10^−3^, see below), and survive or go extinct deterministically because of abiotic and biotic conditions.

We will also vary the dispersal coefficient *D*, to check whether the DL scenario disappears and ET prevails as soon as *D >* 0 or at some higher value of dispersal.

### 1.3 Defining direct and net effects

The matrix *A*_*ij*_ represents *direct effects*, i.e. the instantaneous impact of species *j*’s abundance *N*_*j*_ on the dynamics (per capita growth rate) of species *i* at a given time *t*. By assumption, in the generalized Lotka-Volterra equation (1), these effects are context-independent – they are characteristic of each species pair, and constant across environmental conditions and across the landscape. This provides an important test case for our ability to infer species interactions, since it entails that we can truly assign a ‘ground truth’ value to these interactions that we may hope to recover through some inference method.

However, these direct effects are not necessarily what we try to infer from empirical metacommunity data, where we rarely have access to per capita growth rates. Instead, we typically care about how the presence of a species influences another’s abundance. Since this influence could change over time, we must specify this question further, e.g. ask how a species affects another’s abundance in the long term, at dynamical equilibrium. The roots of this question can be found in early work (Levine, 1976; Lawlor, 1979) who recognized that pairwise species interactions could be separated into direct effects (where species affected each other directly by proximate effects on birth or death rates) and indirect effects (mediated by chains of direct effects from one species to another). This work highlighted that observable effects of species on each other were most often related to ‘net effects’, that involved the entire network of possible indirect and direct effects, and may have little similarity to the direct interactions that drive them.

For mathematical reasons presented briefly here and explained in detail in (Zelnik *et al*., 2024), Levine and later authors proposed that net interactions could be derived as the coefficients in the inverse of the matrix of direct interactions. In the absence of dispersal (*D* = 0), the equilibrium condition for the subset *s*(*x*) of species that coexist at location *x* is given by equation (1) with the content of the parentheses set to 0 (since *dN*_*i*_*/dt* = 0 but *N*_*i*_ *>* 0)

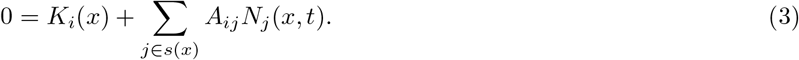

This linear system of equations can therefore be inverted (Levine, 1976) to yield the equilibrium abundances

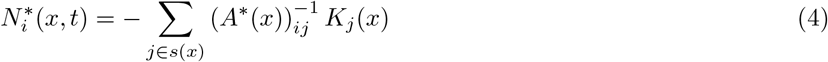

with *A*^*\**^(*x*) the submatrix of *A* restricted to the survivors at site *x*. As a consequence, we can define

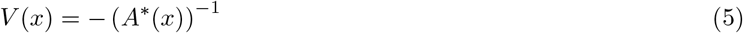

the matrix of *net effects*^1^ at site *x*, which represents the long-term consequences of interactions: how permanently changing the carrying capacity *K*_*j*_ of species *j* (making the local environment more or less favorable to it, e.g. via an experimental treatment) will modify the equilibrium abundance of another species *i*. More broadly, we can define *V*_*ij*_ as how any permanent change in the dynamics (per-capita growth rate) *d* log *N*_*j*_*/dt* modifies the equilibrium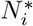.

Clearly net and direct effects are not immediately comparable, since *A* represents the instantaneous impact of one species’ abundance on another’s dynamics, whereas its inverse *V* represents the long-term impact of one species’ dynamics on another’s abundance. Yet it can be shown that, properly defined, net effects can be understood as the sum of all chains of direct effects that connect *i* and *j* via any number of intermediate species in the community Zelnik *et al*. (2024).

### 1.4 Context-dependence of net effects

Schaffer (1981) pointed out that the inverse of a matrix depends sensitively on all its elements, and thus, even for context-independent direct interactions *A*_*ij*_, the specific composition of surviving species can strongly modify the value or nature of net interactions *V*_*ij*_. For instance, adding a third species may cause two competitors to facilitate each other indirectly through their competition with a common enemy. While it may seem counter-intuitive that, say, net effects between two dominant species might change drastically even from adding a rare third species, it is important to point out that these effects are fully realized only in the long term: the inverse matrix appears naturally when computing *equilibrium* abundances (4) so that even a rare or slow-growing species has time to exert or mediate significant impacts on others. Observing abundances out-of-equilibrium could lessen the importance of very indirect paths, and limit context dependence (Zelnik *et al*., 2024).

Clearly, if species composition *s*(*x*) depends on environmental conditions, then so will net effects, even assuming fixed direct effects *A*_*ij*_ across the entire landscape. Fig. 2 demonstrates this context-dependence of net effects.

**Figure 2:**
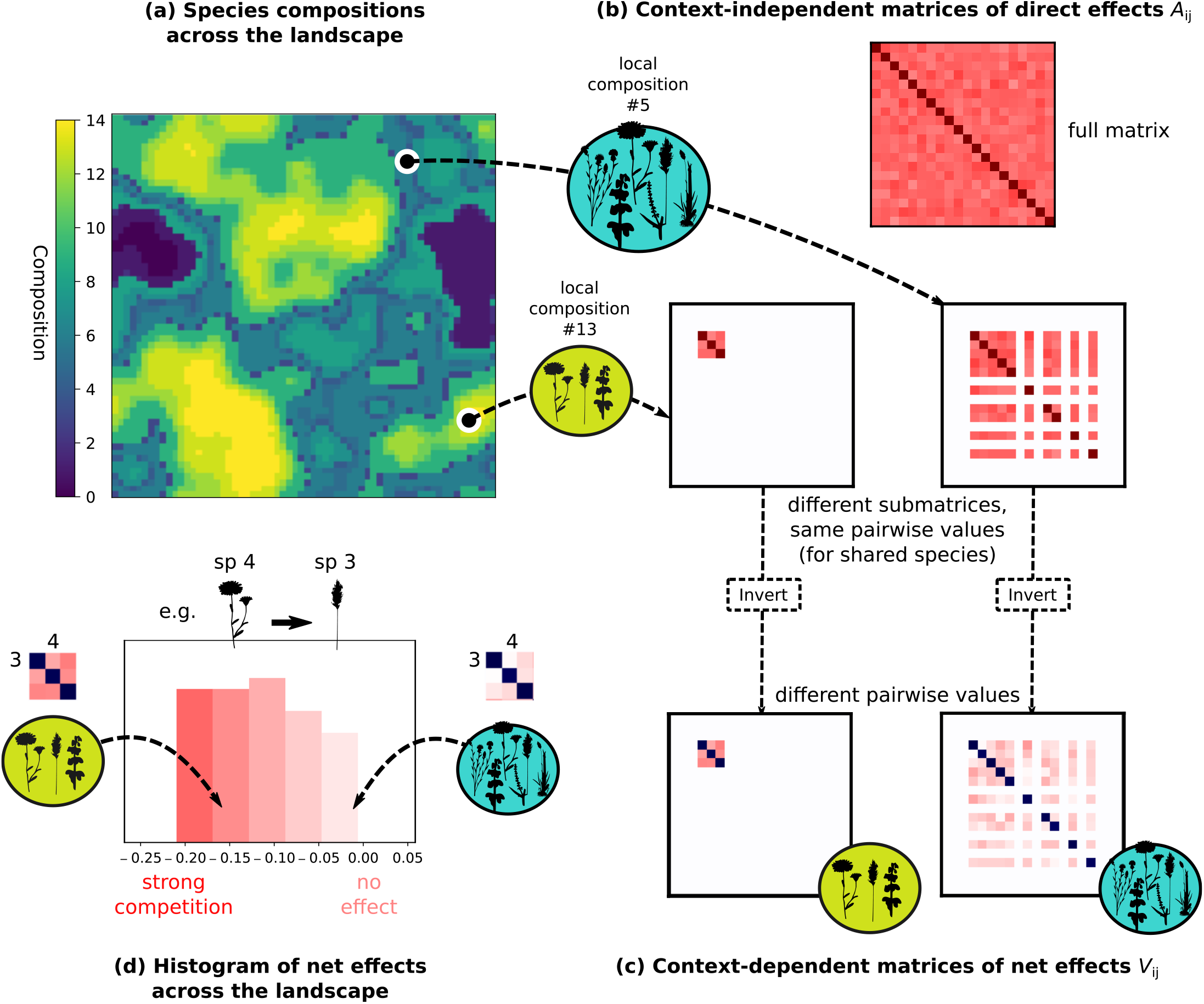
Net effects vary throughout the landscape due to changes in species composition, as illustrated here in one example simulation with dispersal (environment tracking in Fig. 1). While direct effects *A*_*ij*_ (instantaneous impact of species *j* on growth of *i*) are fixed by assumption in our model (1), net effects *V*_*ij*_ (long-term impact of species *j* on equilibrium abundance of *i*) are context-dependent and vary due to the presence or absence of other species. **(a)** A number is assigned to each species composition, and mapped through the landscape. Rare compositions are assigned number 0. **(b)** The fixed matrix of direct effects *A*, and two examples of submatrices restricted to locally surviving species from two different sites. **(c)** Inverting these two submatrices gives local matrices of net effects, *V*_*ij*_(*x*) at each site *x*, where individual elements are now different between localities even for pairs that appear in both localities. **(d)** Histogram of local net effects for the most abundant species pair in the metacommunity (species 3 and 4). The net effect *V*_3,4_(*x*) is computed at every site *x* in the landscape. The mean of these values over the whole landscape gives the spatial average 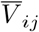 referenced in Fig. 3 and 4.

For non-negligible dispersal *D >* 0, there is no simple linear relationship between abundances and carrying capacities. Therefore, net effects at each site depend not only on local species composition, but also on species abundances in other sites as well as the local abundances. A generalization of *V* near equilibrium can still be made using the inverse of the Jacobian matrix for the full multi-patch dynamics (Gravel *et al*., 2016). Here we always retain very small values of dispersal *D* and this issue does not arise.

### 1.5 Inferring abiotic niches

A frequent objective when modelling species distributions in space is ascertaining the impact of environmental factors, also understood as the ‘fundamental niche’ of a species (i.e. what range of abiotic conditions it tolerates), often seen as a prior step to deciphering the impact of species interactions. It is however understood that biotic interactions transform this fundamental niche into a ‘realized niche’ which can bias our perception of species’ environmental preferences: for instance, certain species might only occur in extreme environments because other competing species prevent them from occupying the more temperate environments that they would prefer (Poggiato *et al*., 2021).

Since our focus is rather on the challenges of inferring biotic interactions even under favorable conditions, we do not delve deeply into the problem of simultaneously determining abiotic niches and interactions, which is central in joint Species Distribution Models e.g. Ovaskainen *et al*. (2017).

Nevertheless, using our simulated data, we could simply try to fit the parameters of equation (2), i.e. the true functional form used to generate the data, ignoring species interactions. This amounts to modelling abiotic niches as carrying capacities *K*_*i*_ that have a Gaussian dependence in the environmental factor *E*, with species-specific optima and widths (Fig. 1). As we show in Appendix, within the dynamical regime and parameter range considered in our simulations, species interactions are indeed distorting the apparent relation between abundance and environment (Fig. S1 and S2).

Yet this distortion remains sufficiently limited in our case that an observer would get a passable estimate of each species’ environment preferences, i.e. the center of its fundamental niche and a lower bound on its width, simply by fitting a Gaussian curve to the maximum abundance seen in each environmental condition (details in Appendix).

Therefore, in the rest of the main text, we entirely bypass the issue of inferring abiotic niche, and investigate inference challenges that remain even assuming that we perfectly know the carrying capacities of every species at every location.

### 1.6 Inferring biotic interactions

We infer biotic interaction effects through multilinear least squares regression in two distinct ways. On one hand, we can infer estimates of **net** effects 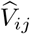 as the multilinear coefficients in

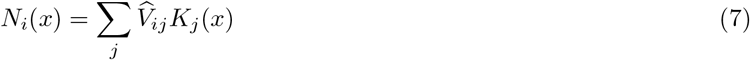

On the other hand, following (Xiao *et al*., 2017; Barbier *et al*., 2021), we can infer **direct** effects *Â*_*ij*_ as the multilinear coefficients in

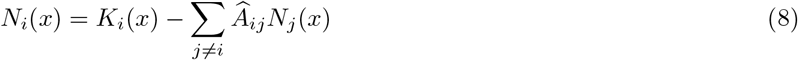

In the first case, we need to know the abiotic niches, i.e. carrying capacities *K*_*i*_(*x*) across the landscape. In the second, we can either use known carrying capacities, or infer them as the (site- or environment-dependent) intercept of the relation between the abundance of species *i* and other species.

Since we noted above that deducing environmental preferences is not the most severe obstacle in our chosen simulation setting, we hereafter assume that carrying capacities *K*_*i*_(*x*) determined by the abiotic factor *E*(*x*) are known prior to inferring interactions (e.g. if they are measured in experiments or can independently be estimated from species distributions). We aim to show that even this favorable case presents considerable difficulties, that exist independently from the problem of inferring environmental effects (see Appendix for further discussion).

## 2 Results

### 2.1 Direct and net effects

Modern inference methods, e.g. (Ovaskainen *et al*., 2017), attempt to simultaneously deduce species’ interactions, environmental preferences and migration effects from noisy or limited data. Yet significant methodological or conceptual difficulties may arise even with less ambitious goals. Here, we mainly discuss the possibility of estimating species interactions under optimal data conditions: sampling the whole landscape at equilibrium, without any measurement error, and having full knowledge of carrying capacities *K*_*i*_(*x*) (i.e. environmental preferences) for each species.

We break down this issue into the inference of *direct* effects and *net* effects whose definitions we recall here (see Methods for more details). Direct effects correspond to the matrix *A*_*ij*_ in (1) which describes how the current abundance of species *j* influences the instantaneous dynamics (growth or mortality rate) of species *i*. Net effects are given by the matrix *V*_*ij*_ in (5) which describes how a permanent change in species *j*’s dynamics (e.g. a change of carrying capacity, or removal from the community) would impact the abundance of species *i* in the long term Zelnik *et al*. (2024). Direct effects are not usually what we try to infer in a biogeographic context since we rarely have access to the population growth rates, but they mediate and explain net effects on abundances.

We see in Fig. 3(a,c) that the inference of *direct* effects still depends on the colonization scenario: it is successful with dispersal limitation (DL), but not with environment tracking (ET). In the DL scenario, the inferred matrix *Â*_*ij*_ is very similar to the ground truth matrix *A*_*ij*_, missing only a few interactions for species that are never present together in the landscape. In the ET scenario, estimates *Â*_*ij*_ are usually wrong and may even have the wrong sign, though the inference tends to improve here for stronger interactions (as this leads to fewer species coexisting, and thus a simpler inference task).

**Figure 3:**
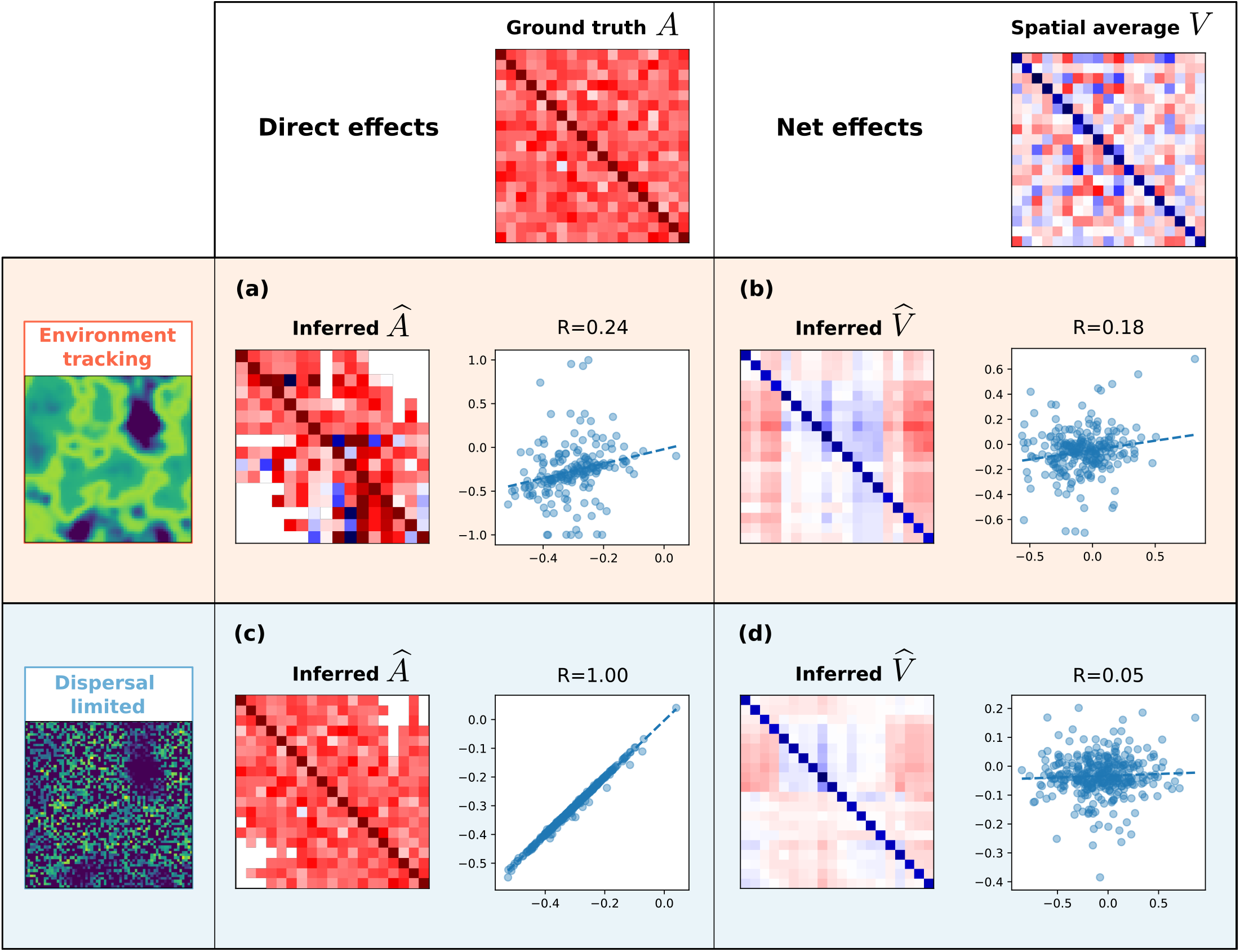
Inferring the full matrix of direct or net effects in the example simulation run under optimal conditions (noiseless data, full knowledge of the abiotic niche of each species, no dispersal, see Methods). Each point corresponds to a pairwise effect of species *j* on species *i*. Dashed lines indicate linear regressions. Each row corresponds to a colonization scenario (see Fig. 1 and Sec. 1.2). **(a**,**c)** Direct effects. On the y-axis, values *A*_*ij*_ inferred through hyperplane regression of *N*_*i*_(*x*) against *N*_*j*_(*x*); on the x-axis, ground truth matrix, *A*_*ij*_. **(b**,**d)** Net effects *V*_*ij*_. On the y-axis, values 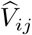 inferred through hyperplane regression of *N*_*i*_(*x*) against *K*_*j*_(*x*). On the x-axis, “ground truth” obtained by spatial average over local matrices, 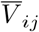(see Methods).

As for net effects, in our Lotka-Volterra model they are ill-defined at the landscape scale: the value *V*_*ij*_ depends on species composition, i.e. the set of surviving species, which varies across the landscape (Fig. 2). Still, we could hope that the *average* value ⟨*V*_*ij*_⟩ across the landscape (Fig. 2c) is meaningful. In that case, we expect it should correlate with the apparent net effect, defined as the regression slope of *N*_*i*_ against *K*_*j*_ (putting together values measured across the whole landscape).

We find in Fig. 3(b,d) that this is not the case, which can be explained by the fact that the value of *V*_*ij*_ and *K*_*j*_ are actually very correlated across the landscape, so a spatial average of *V*_*ij*_ is not representative of local net effects (Fig. 2). We conclude that inferring landscape-scale net effects is an ill-posed problem, as they are not well defined even when direct effects are assumed constant.

### 2.2 Interaction statistics

While the full matrix inference is unsuccessful in many cases, we see in Fig. 4 that interaction statistics are more robustly estimated. Indeed, there is a very strong correlation between the ground truth mean interaction strength ⟨*A*⟩, and the mean measured over our empirical estimates ⟨*Â*⟩, even when the individual elements *Â*_*ij*_ are unsuccessfully estimated.

**Figure 4:**
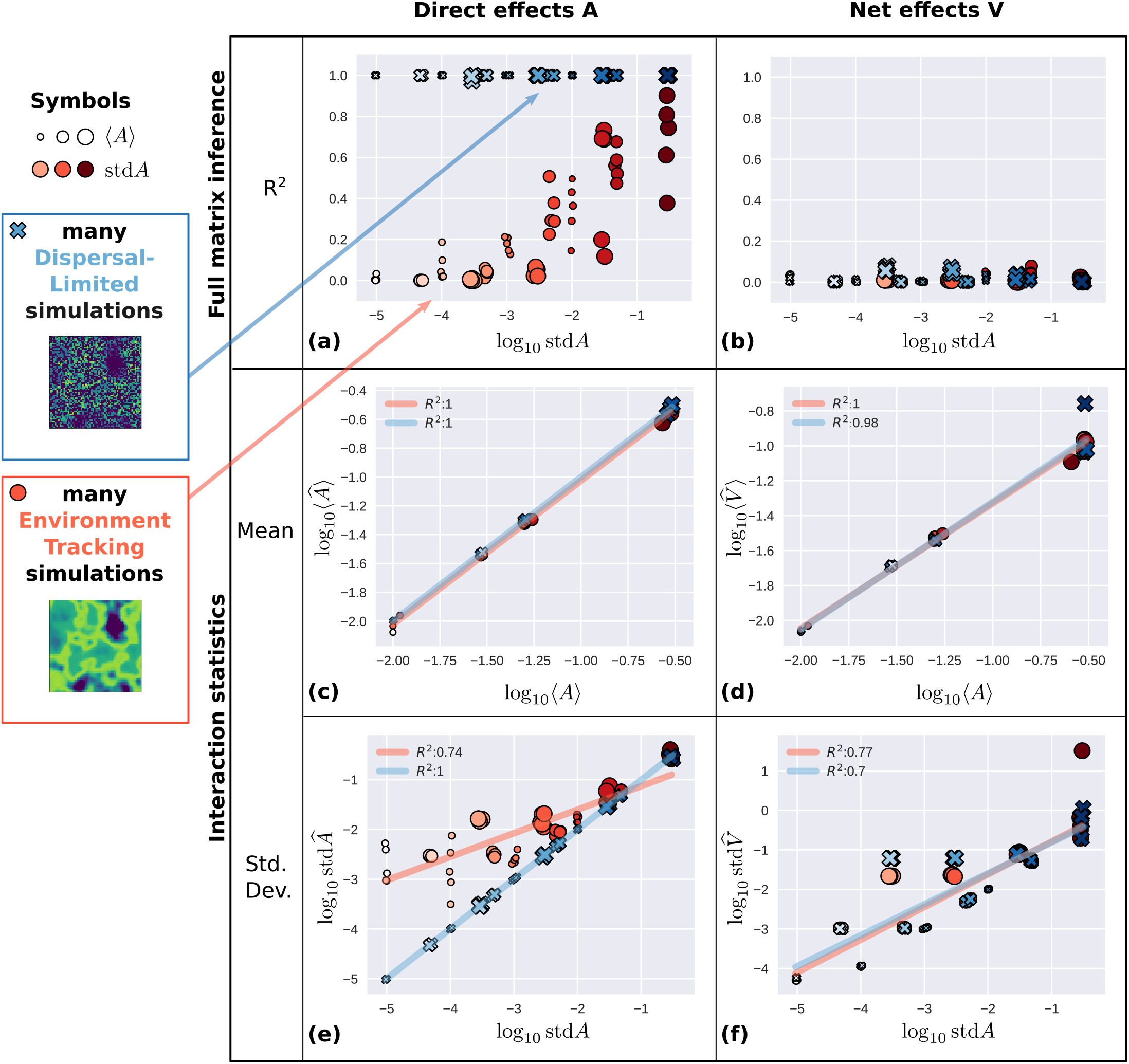
Inferring all interactions or only statistics across many simulation runs with different parameters. Each point is a simulation run, corresponding to one of two colonization scenarios (ET: orange circles, DL: blue crosses) and various values of ground truth statistics ⟨*A*⟩ ∈ [0.01, 0.03, 0.05, 0.3] (symbol size) and std(*A*) (symbol color saturation, obtained by multiplying each value of ⟨*A*⟩ by a value in [0.001, 0.01, 0.1, 1]). **(a**,**b)** *R*^2^ of full matrix inference (see e.g. Fig. 3 for one simulation run) is only successful for direct effects under dispersal limitation. However, the statistics of inferred interactions are robustly related to ground truth statistics, for both inferred direct and net effects, *Â*_*ij*_ and 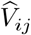. This relationship is very strong for mean interaction **(c**,**d)**, and weaker for standard deviation **(e**,**f)**.

Likewise, there is a strong relationship between ⟨*A*⟩ and average net effects across the landscape, ⟨*V*⟩ (Fig. 4d). These two quantities are not equal even in principle, but a robust relationship suggests that we could use one to infer the other. Standard deviations are also correlated between ground truth and inferred values, though more weakly (Fig. 4e,f). We notice that they are more sensitive to the colonization scenario (DL or ET) and thus we can only roughly deduce the true std(*A*) from its empirical estimate (especially at small values, std*A <* 10^−2^) without knowing which dispersal scenario we are observing. Finally, symmetry is perfectly estimated in the DL scenario, but entirely incorrect in the ET scenario (see Appendix, Fig. S4).

To summarize, it may be possible to infer the full matrix of direct effects for abundant data, with dispersal limitation or some other phenomenon decoupling species composition from the environment. The full matrix of net effects is not well-defined, and no inference method is successful. Under broader conditions, we can likely only estimate statistical features, most reliably the mean interaction strength.

### 2.3 Influence of dispersal

We show in Fig. 5 that, as we increase dispersal coefficient *D* from zero, the transition between dispersal limitation (DL) and environment tracking (ET) scenarios is abrupt in our model, occurring for *D ≈* 10^−3^.

**Figure 5:**
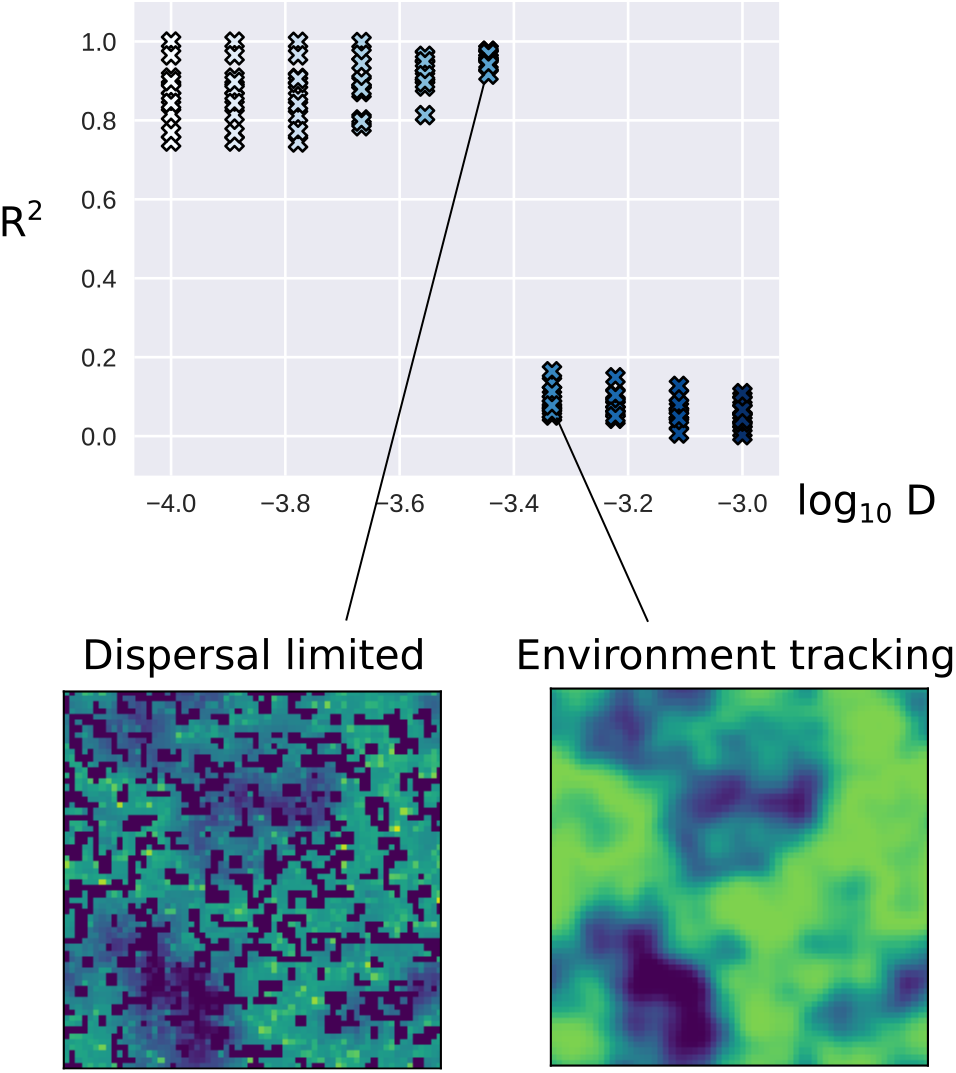
Effect of dispersal on the ability to infer the full matrix of direct species interactions *A*_*ij*_. On the x-axis we represent the dispersal coefficient *D* in log 10 scale. On the y-axis we show the Pearson *R*^2^ of the fit between true and inferred net effects. Each symbol is a simulation out of sets (all sets share the same landscape, each set has a distinct interaction matrix, and each simulation within a set differs by its value of dispersal). Below, we show the final abundance of one species across the landscape in one simulation set, for values of dispersal right below and above the transition from dispersal limitation to environment tracking. The patchy appearance of the left-hand inset is due to migrant fluxes from neighboring patches being too weak in many cases to overcome the local extinction threshold, as explained in Section 2.3.

When dispersal becomes able to overcome the initial absence of a species in a patch, by creating a migrant flux above the extinction threshold *N*_*c*_ and thus allowing species that can invade the patch to reach a nonzero equilibrium, local species composition becomes entirely determined by each site’s biotic and abiotic environment (rather than initial conditions and dispersal) and the ability to separate the influence of various species decreases abruptly, as their abundances co-vary much more strongly (which may be seen in Fig. S5 in Appendix).

The abruptness of the transition is due to the fact that our model has a sharp extinction threshold *N*_*c*_ = 10^−3^. Thus, patches where dispersal cannot bring the abundance above the threshold cannot be colonized by the species. Many different species compositions in neighboring patches are preserved until dispersal allows crossing this threshold systematically for all species. However the total number of observed species compositions across the landscape is *not* sharp in *D* (Fig. S6 in Appendix), suggesting that the ability to correctly infer detailed interactions is not tied to the diversity of compositions over the whole landscape, but rather simply to the existence of diverse compositions in close proximity.

## 3 Discussion

Many statistical and theoretical methods have been proposed to infer ecological interactions between species from their spatial co-distributions. The best-studied obstacle to this inference is the possible confounding effect of other factors that impact spatial distributions, e.g. the fact that species may appear positively associated because they have similar environmental preferences (Ovaskainen *et al*., 2017). Recent studies have discussed issues with methods devised to overcome this obstacle, for instance, that interactions may prevent us from correctly understanding species’ environmental preferences (Poggiato *et al*., 2021) Here, we have mainly focused on two further obstacles to the precise inference of species interactions, arising even when we are in the most favorable conditions to address the problems noted previously.

### 3.1 Statistical issues and non-identifiability

One obstacle is statistical in nature, i.e. difficulties in identifying the model due to multi-colinearity. We find that we can successfully infer direct species interactions only in scenarios where species composition is forced to vary substantially and independently from the environment, for instance due to dispersal limitation or experimental manipulation (as in biodiversity experiments). We cannot do so if the same environmental conditions predictably lead to the same species composition, a situation that we call “environment tracking”.

The issue is not only that effects of interacting with particular species may be confounded by environmental effects, but they may also be confounded by each other, as the abundances of multiple interaction partners tend to covary positively or negatively based on their similar or dissimilar environmental preferences. That latter problem decreases when interactions have more variance (Fig. 4a) or when species are differently influenced by many independent environmental factors (see Appendix, Fig. S7), but this does not suffice here to completely eliminate the problem of model identifiability.

All our results were obtained in a setting that should be highly favorable to the detection and estimation of species interactions: there are context-independent parameters defining direct interactions, species often coexist, they reach stable abundances that reflect their preferences, dispersal is small and intervenes mainly to allow species to colonize patches where they were not seeded initially (but does not significantly distort equilibrium abundances). Despite all these favorable assumptions, we found that it may not always be possible to precisely infer the details of species interactions. Relaxing some of these assumptions to consider stronger or more context-dependent interactions (e.g. priority effects, environmental modification), more complex dynamics (chaos, transients, external perturbations), stronger spatial fluxes, and observational issues (data limitation, errors and biases), is likely to introduce further difficulties but perhaps also different opportunities for the inference of interactions. Indeed, we speculate that various obstacles could work against each other: stronger non-linearities may somewhat alleviate model non-identifiability; conversely, having a large number of species and parameters may end up being the main challenge in model fitting and override the importance of details of how each interaction is modelled; finally, other dynamical settings, such as species abundances fluctuating chaotically rather than being at equilibrium, may require entirely distinct methods with different challenges.

A more conceptual problem lies in the context-dependence of interactions: there might not exist any constant number that would adequately represent “the effect of species *j* on species *i*” across a whole metacommunity, in which case our inference problem is ill-posed from the start. On the one hand, the Lotka-Volterra model used here (1) can be thought of as giving a lower bound on the amount of context-dependence we can expect in a plausible ecological setting. The model assumes total context-independence of all direct effects (per-capita instantaneous impacts on growth rates) between species. Yet, the ‘net effects’ between species, defined to include all indirect impacts arising over time and through intermediates (e.g. indirectly helping a species by directly hindering its competitors), are found to be highly context-dependent as soon as interaction strength is not very small (Zelnik *et al*., 2024). On the other hand, our choice of looking at long-term abundance patterns, letting species reach an equilibrium, is giving maximal opportunity for such indirect effects to play out – even a rare or slow-growing species has time to exert or mediate significant impacts on others, hence the fact that the matrix of net effects can change drastically when we remove a species, no matter how rare. Thus it may be that observing abundances out of equilibrium, driven by more complex ecological dynamics or external perturbations, and tracking temporal (or spatio-temporal) rather than purely spatial co-distributions, could lessen the interference that might be due to this context dependence and entanglement arising in communities at equilibrium.

Our work stresses the importance of correctly specifying which concept of biotic interaction one is trying to infer: for instance, estimating context-independent direct effects is sometimes possible even when net effects vary dramatically across the landscape. This is of particular relevance to statistical approaches focusing on the co-distribution of species, e.g. joint Species Distribution Models Ovaskainen *et al*. (2017). The residual covariance between species across the whole landscape can be understood as the spatial aggregation of locally varying net effects which we believe (see Appendix: Estimating interactions from residual species co-variation) is not an appropriate path to deduce direct effects.

### 3.2 Getting more from less

Despite these two obstacles, we also found that the community-wide statistical properties, i.e. mean and variance, of direct interaction coefficients could be relatively well inferred from the observed species distribution patterns, even when our detailed estimates of pairwise interactions were entirely incorrect. We did not attempt here to develop novel techniques specifically for the purpose of inferring moments of the distribution of interaction strengths. Instead, we used simple methods to estimate all pairwise interactions, and then computed the moments of these estimates, even when they were individually wrong. It is likely that methods that would be tailored to capture statistical moments directly would be even more robust. But it is rather striking that applying the ‘wrong’ method still provides reasonable estimates of the moments: it suggests that, even when observational or experimental attempts at measuring interaction strengths (e.g. (Barbier *et al*., 2021)) yield incorrect numbers, these numbers might still have the right statistics to characterize how important and diverse species interactions are overall in the community-level ecological dynamics.

From an empirical point of view, our results thus carry encouraging as well as cautionary notes. Species-level questions may require the precise inference of a given pairwise interaction, and our work comes here as a warning to keep in mind possible barriers to that inference when we do not have grounds to assume ‘natural experiments’ such as permitted here by strong dispersal limitation. On the other hand, we feel that for empiricists interested in estimating the intensity of biotic interactions in a community as a whole, e.g. to know whether community composition is a better indicator of environmental states or internal dynamics, our findings are encouraging and a suggestion to turn to methods that strive to estimate community-wide statistics rather than individual pairwise species interactions.

Finally, we must consider the empirical relevance of our study’s assumption that direct species effects are context-independent and simply additive with each other and with environmental effects. This may seem like a highly restrictive assumption, limiting the value of trying to infer such direct effects. We nevertheless speculate that two facts make this assumption less restrictive than it seems: first, additivity is more likely to hold approximately in direct effects, which occur over a short time, than in long-term net effects; and second, the congruence of many causal factors (species, environmental variables) hopefully means that the details of each and how they interact matter less, and additive effects may be a mechanistically wrong but phenomenologically useful abstraction, at least when focusing on community-wide statistics and outcomes as we are suggesting here.

### 3.3 Conclusions

In conclusion, metacommunity ecology provides a more comprehensive conceptual framework than the approaches that set the stage for inferring species interactions from co-distributions (Diamond, 1975; Connor & Simberloff, 1979). Work to date relating metacommunity ecology to species co-distribution patterns (Morueta-Holme *et al*., 2016; Ovaskainen *et al*., 2017; Leibold *et al*., 2022; Christopher D. Terry *et al*., 2023) is providing some exciting new tools, but the connection between distribution patterns and ecological mechanisms remains elusive. This is, in large part, because correlations between species are highly sensitive, entangling multiple ecological processes and potentially the entire biota, as put forward by Schaffer (1981).

We find that progress might be made by focusing on less detailed and more robust descriptions of distribution patterns. While it may rarely be possible to infer the full set of parameters describing a metacommunity, it might be more feasible to parameterize models inspired by statistical mechanics, e.g. (Gravel *et al*., 2016; Bunin, 2017), in which only overall statistics of parameters are used to predict a variety of ecological outcomes such as abundance distributions, dynamics and stability. In these so-called “disordered systems” models, outcomes are robust to changing many details of interaction coefficients, as long as species are not organized in a well-ordered structure such as a strict competitive hierarchy. Further refinements of these models have been proposed to capture important large-scale features of the ecological structure of interactions (Barbier *et al*., 2018). First steps toward parameterizing such models from empirical species co-distributions are being taken in very recent studies (Camacho-Mateu *et al*., 2023) and we expect that continued progress along these lines will prove timely and useful.

## Code availability

All simulation code is written in Python and available from the following repository: https://github.com/mrcbarbier/morefromless

## Acknowledgments

We would like to thank Jean-François Arnoldi, Virginie Ravigné, Kara Taylor, Veronica Franz, Jeff Mintz, and Chase Rakowski for discussions during the elaboration of this study and providing useful comments on earlier drafts. M.B. was supported by the CIRAD funding CRESI 2022. G. B. was supported by the Israel Science Foundation (ISF) Grant No. 773/18. M.A.L. was supported by NSF 2025118 and NSF 2224331 grants.

## Supplementary Materials

### A Inferring carrying capacities

We compare in Fig. S1 and Fig. S2 the fundamental abiotic niche *K*_*i*_(*E*) for each species to the observed distribution of species abundances, to see whether the former can easily be deduced from the latter or appears distorted due to biotic interactions.

The principle we use here is very simple: since interactions are almost all competitive in most of our simulations (some of the randomly-drawn values may be positive, representing mutualism, commensalism, etc. but they are infrequent), we expect the abundance to be at most equal to *K*_*i*_(*E*) most of the time. Therefore we devise a very simple estimate of the fundamental niche by (1) binning observed species abundances into 20 bins based on the value of the environmental factor, (2) computing a higher bound within each bin (using the 95% percentile to be more robust to outliers), then (3) fitting equation (2) with *μ*_*i*_ = 0, i.e. a Gaussian curve, to these local maxima by linear least squares regression, species by species.

(We note that in our simulation model, *K*_*i*_ can be negative, see (2), while the inference approach we use here can never correctly ascertain how bad an environment is when the species does not grow in it, so we do not attempt to fit the ‘mortality’ *μ*_*i*_ defined in (2))

In Fig. S1, we see that the Dispersal Limitation scenario may help to correctly deduce the fundamental niche since there are many different species compositions in the same abiotic conditions, and some of these compositions may be closer to the species being alone, thus we generally recover the overall shape of the niche, though its precise parameters are slightly biased.

In Fig. S2, we observe that in an Environment Tracking scenario, the shape of the realized niche may be quite different from the fundamental niche, as the (mainly competitive) interactions may restrict species to only a small part of their potential range. The inference of niche parameters is not as good; still, the environmental gradient is strong enough here that we can get a rough estimate of abiotic properties through our simple inference method. Given that the ET scenario is in any case unfavorable to inferring species interactions, we do not discuss the additional complexity of having wrong estimates of the carrying capacities *K*_*i*_, and treat them as exactly known when trying to estimate interactions, but this additional difficulty could be discussed in future work.

We show in Fig. S3 how errors made on the inference of each aspect of the niche vary continuously with the dispersal coefficient, interpolating between the two extreme scenarios shown in the previous two figures.

### B Inferring symmetry

Beyond interaction mean and variance, interaction symmetry has often been proposed theoretically as an important parameter, see e.g. (Bunin, 2017; Barbier *et al*., 2018). That was another statistical parameter that we varied and attempted to infer, see Fig. S4. We find that interaction symmetry is only correctly inferred for direct effects, and only when the full matrix inference is successful, i.e. in the Dispersal Limited scenario.

### C Estimating interactions from residual species co-variation

Consider patterns of covariation across space:

- Covariance of carrying capacities “cov*K*”:

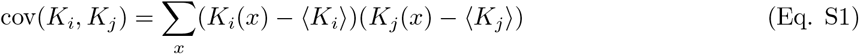
- Covariance of abundances “cov*N* “:

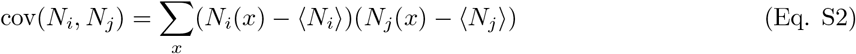

To see whether a trace of biotic interactions can be found in residual covariation, we first decompose the covariance of abundances into two components:

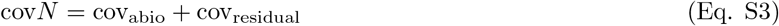

The abiotic component is simply derived from the covariance of carrying capacities

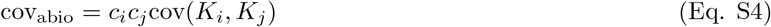

with unknown scaling coefficients *c*_*i*_ that are inferred by least squares regression.

This mimics the way that a simple SDM would predict species covariation (for each species *i*, a regression between abundance *N* and the environmental factor). Given (5), we can expect that in the case of weak net effects between species, this formula is approximately correct for our dynamical model with *c*_*i*_ *≈ V*_*ii*_.

We can then study whether cov_residual_ contains information about the full matrix or statistics of direct effects *A*_*ij*_ or net effects *V*_*ij*_.

All the inferences in the main text were performed assuming that we have full knowledge of abundance *N*_*i*_(*x*) and carrying capacity *K*_*i*_(*x*) for every species *i* at every point in space *x*. Here we assume that we have access to less information: only how abundances or carrying capacities covary between species across the landscape, shown in Fig. S5(a,b,c).

Removing the covariation between species due to the environment, we can observe how statistics of the coefficients *c*_*i*_ in the regression of abundances to the abiotic niche *N*_*i*_ *∼ c*_*i*_*K*_*i*_(*E*(*x*)), and residual covariation cov_residual_ depend on the mean and variance of interactions. We see in Fig. S5 that there is no clear relationship with ground truth parameters. It thus seems impossible, under the studied model setting and approach, to infer even only interaction statistics without having more detailed information on abundances and carrying capacities.

**Figure S1:**
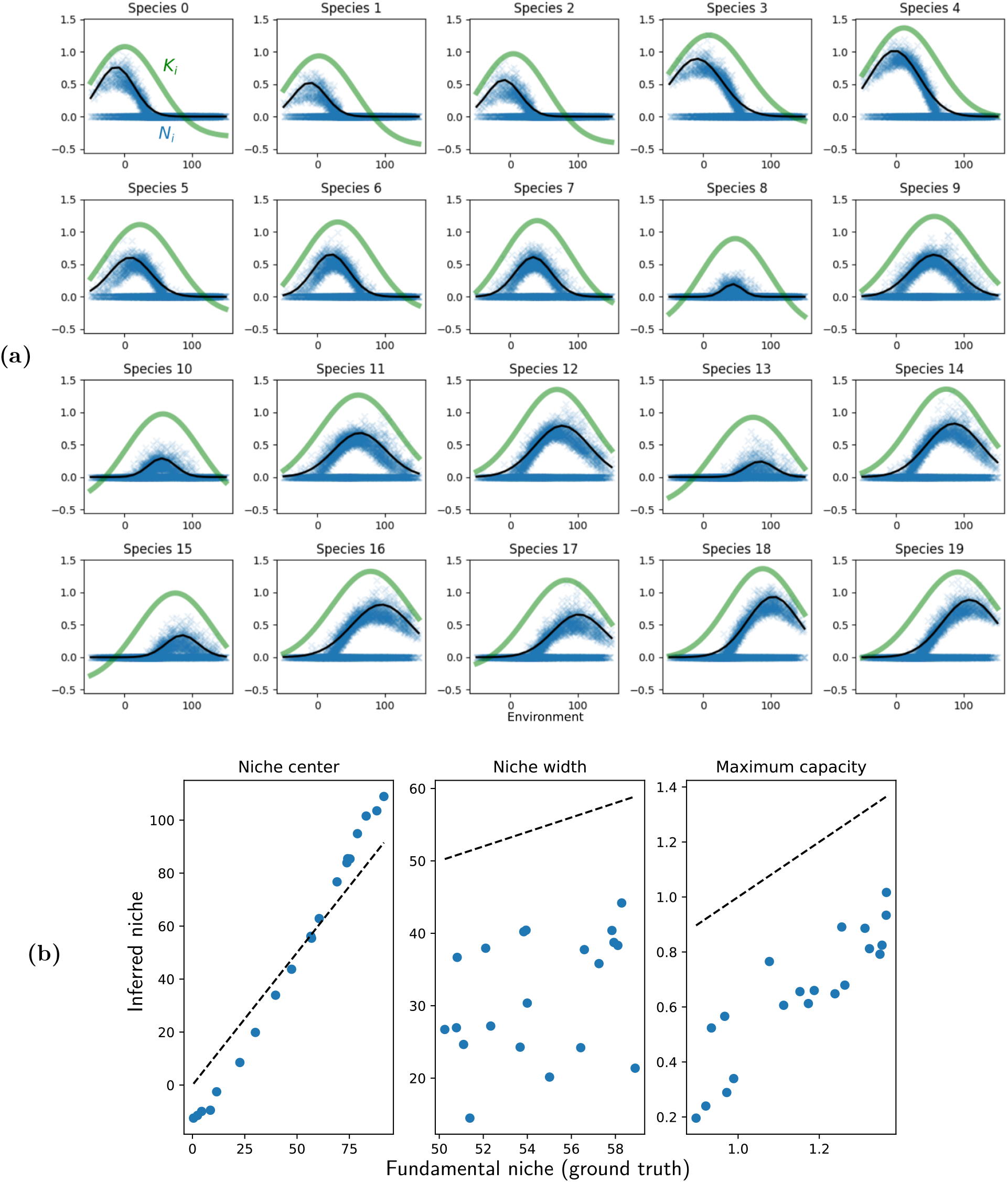
Inferring carrying capacities in one simulation example under conditions of dispersal limitation, see details in Appendix A. **(a)** Fundamental and inferred niches for each species, as a function of values of the environmental factor *E ∈* [–50, 150]. Solid green curve: fundamental niche (carrying capacity as a function of environment). Blue symbols: abundances observed throughout the landscape. Solid black curve: niche inferred through our method. **(b)** Comparing the center, width and maximum of the fundamental (x-axis) and inferred (y-axis) niches.

**Figure S2:**
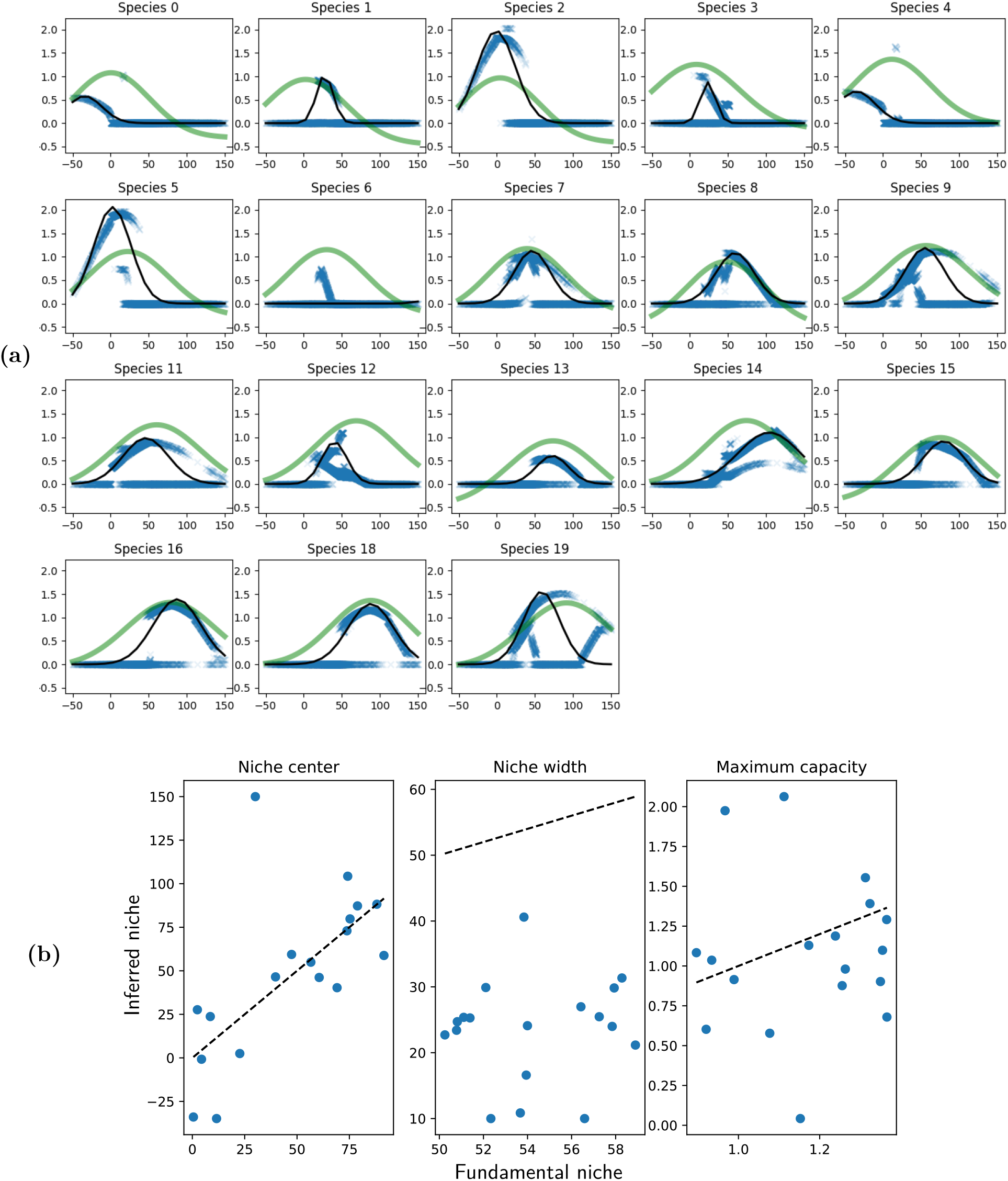
Inferring carrying capacities in one simulation example under conditions of environment tracking, see details in Appendix A and Fig. S1. Some species have gone extinct and are not represented in (a).

**Figure S3:**
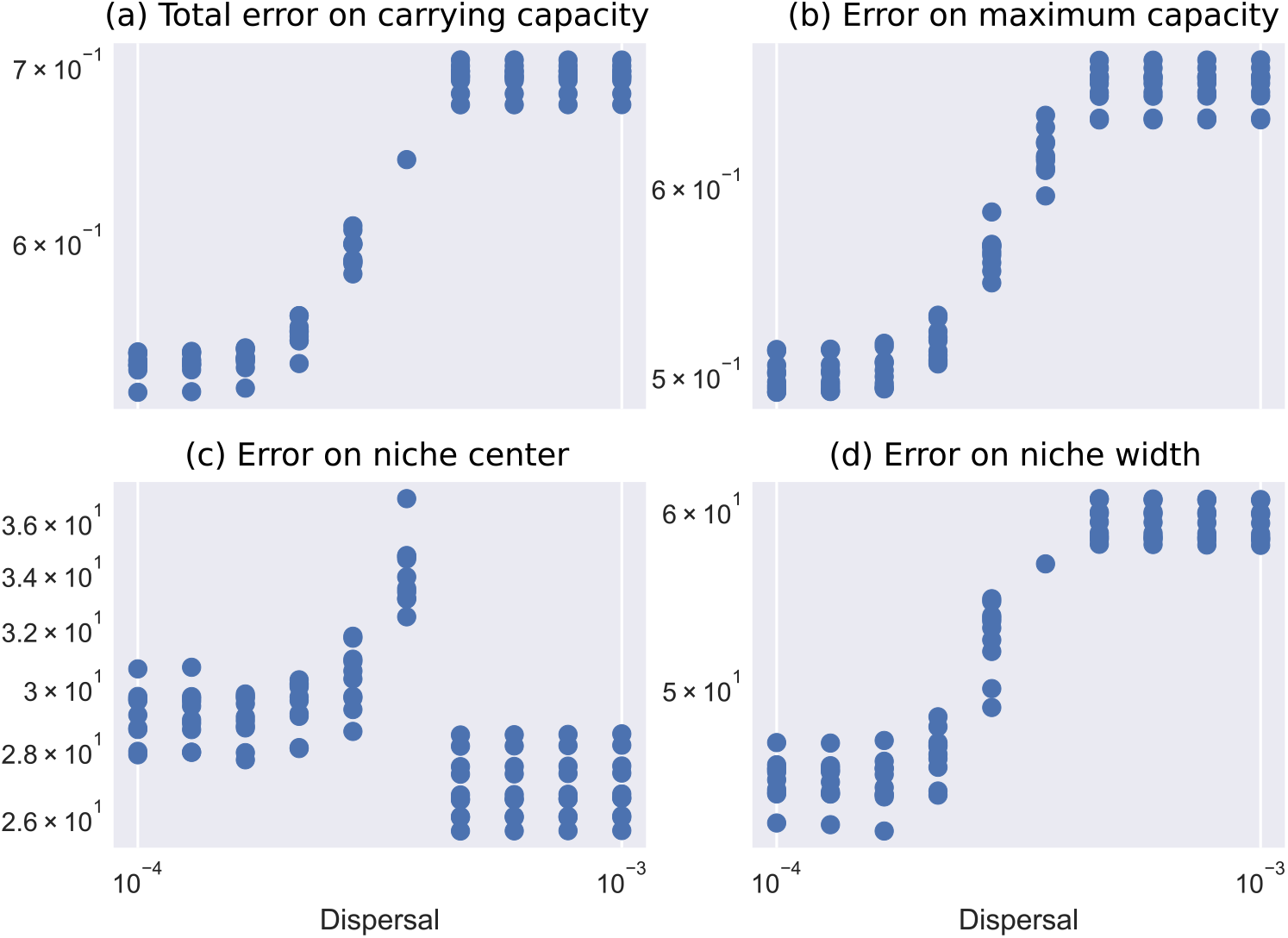
Impact of dispersal coefficient *D* on error on **(a)** estimates of carrying capacity at each point of the environmental gradient, **(b)** estimates of maximum carrying capacity, **(c)** niche centers, **(d)** niche width. In each case, errors are computed as the square root of the average across species – and average across space in case (a) – of (groundtruth value - inferred value)^2^, with the inference approach discussed in Appendix.

**Figure S4:**
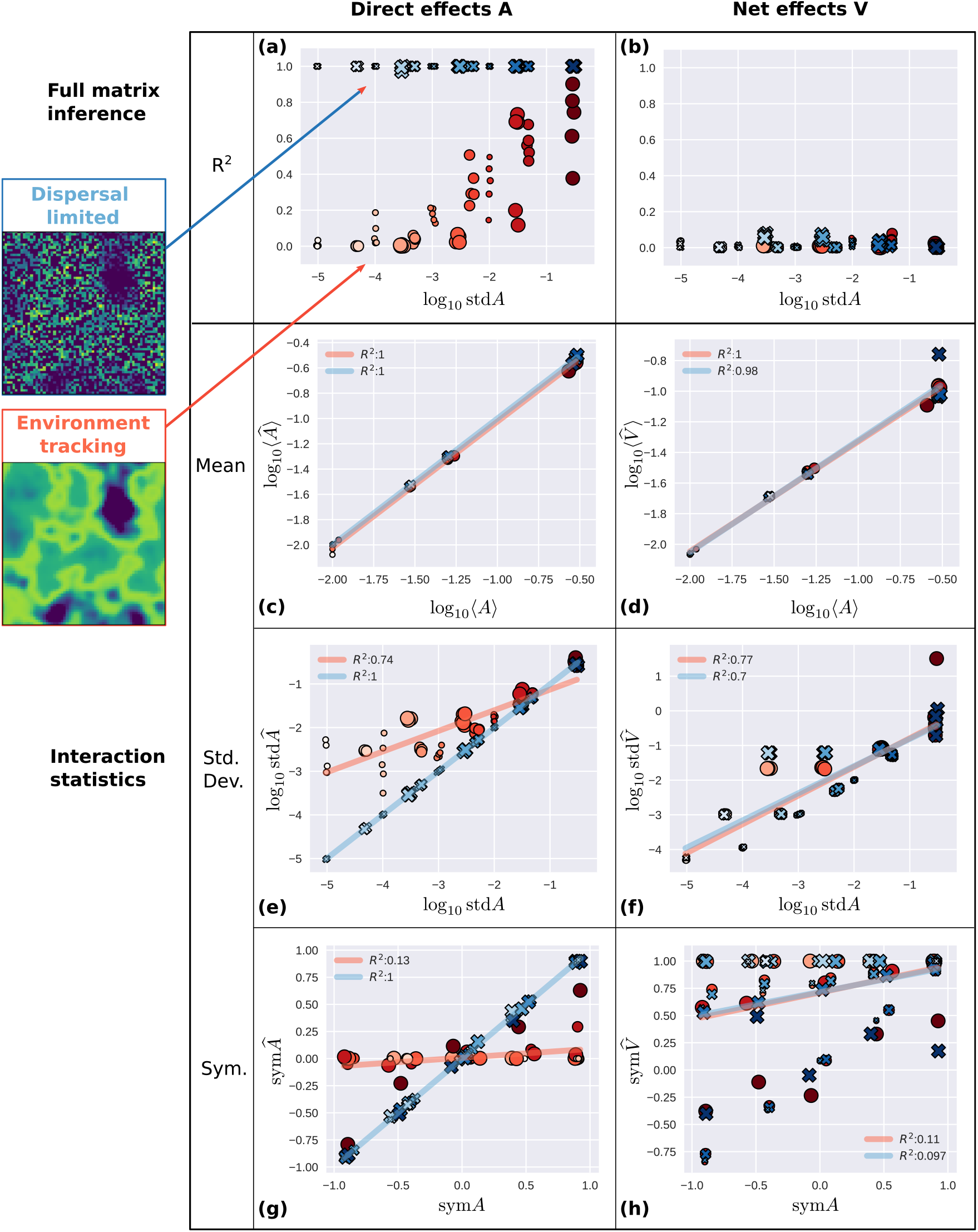
Extension of Fig. 4 including interaction symmetry **(g**,**h)** which is only correctly inferred for direct effects, and only when the full matrix inference is successful, i.e. in the DL scenario.

**Figure S5:**
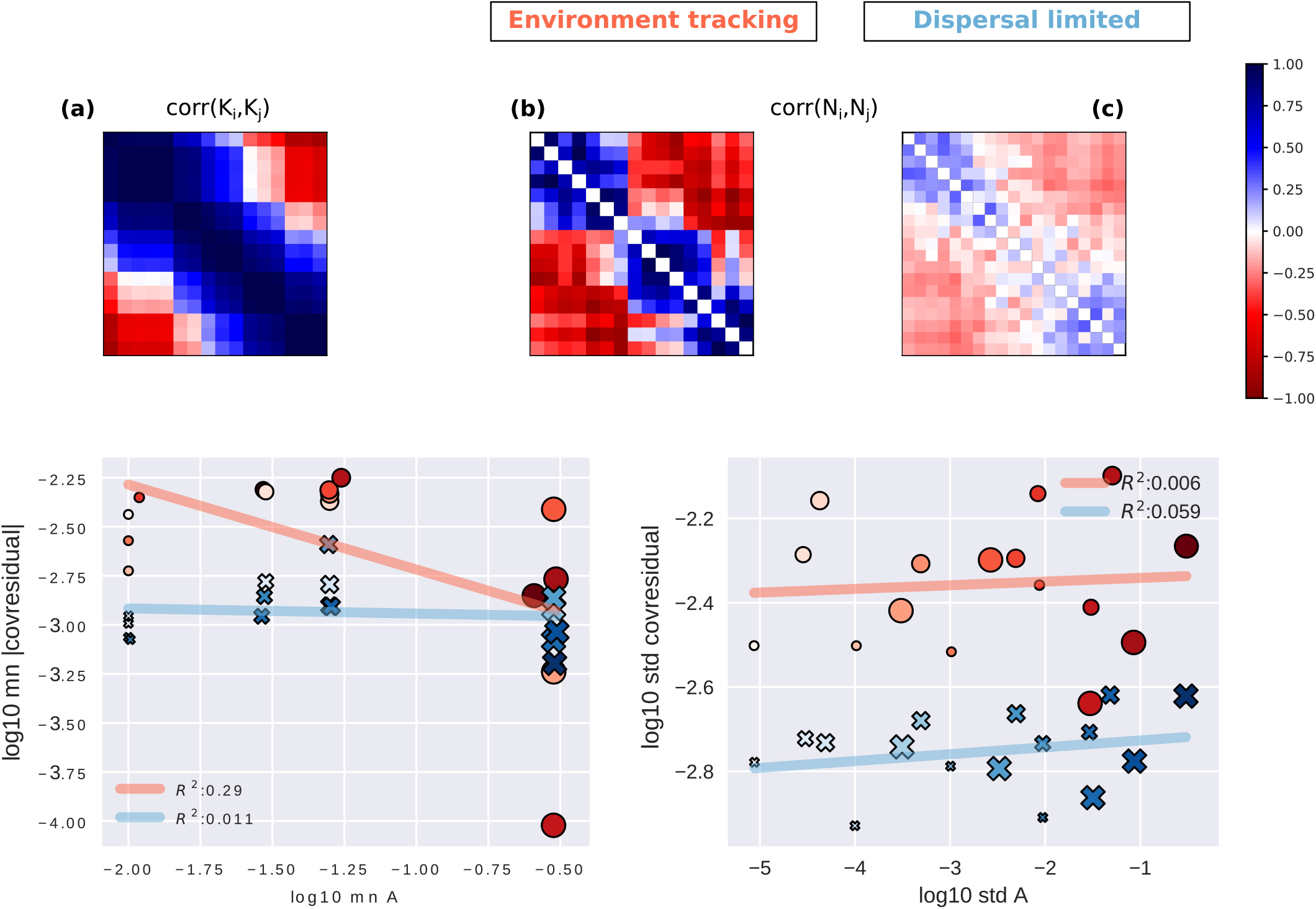
Trying to infer interaction statistics using only the knowledge of covariance or correlation of (a) K and (b-c) N in the two scenarios. From these observations, we compute the residual covariation of species once we control for environment-driven covariation (see Appendix C), and compute its statistics: (d) mean and (e) std. We see no relationship between the statistics of interactions and residual covariation, suggesting that this limited knowledge is insufficient to correctly detect interaction strength

**Figure S6:**
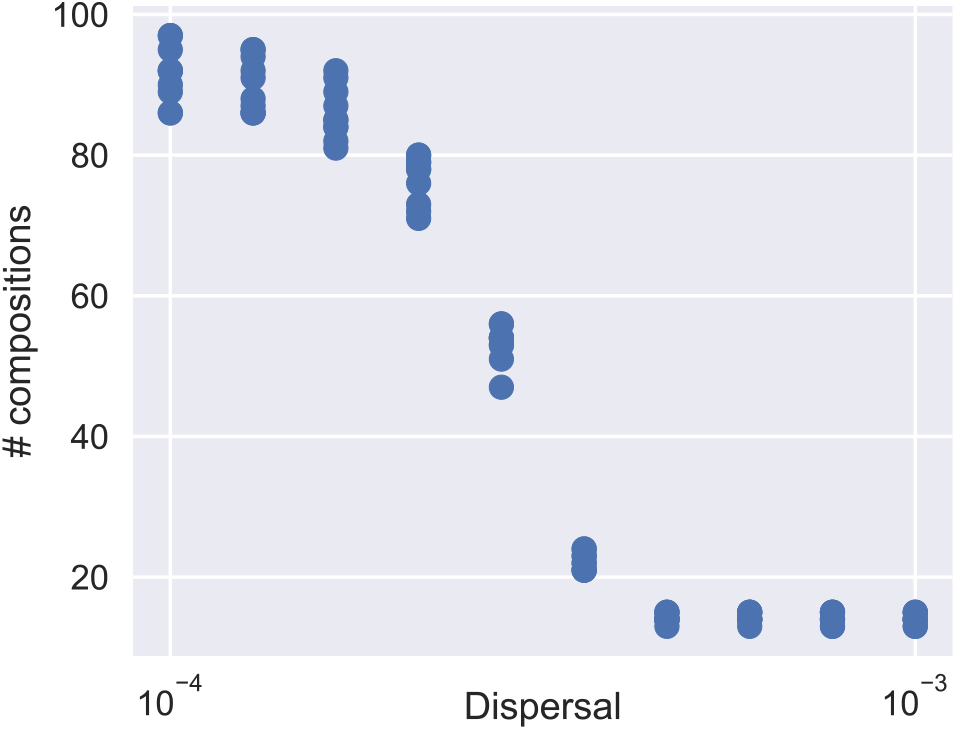
Number of compositions versus dispersal coefficient, showing no abrupt transition contrary to the success of interaction inference in Fig. S6.

**Figure S7:**
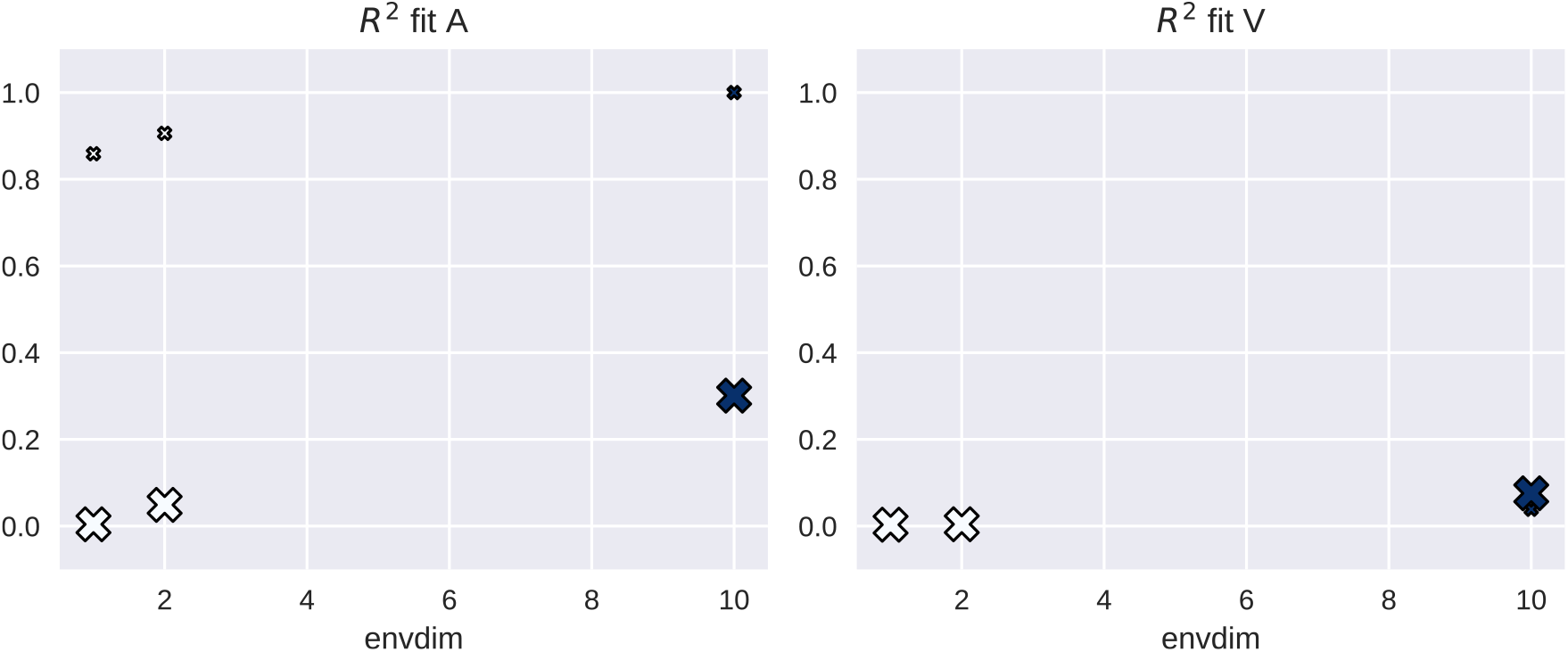
*R*^2^ of inference of direct and net interactions (*A* and *V* respectively) depending on the number of independent environmental factors varying through the landscape (1, 2 or 10, shown on the x-axis), and depending on dispersal (small symbols: *D* = 0, large symbols: *D* = 0.3). When there are multiple environmental factors *E*_*j*_, equation (2) is amended to become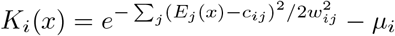.

## Footnotes

1 We note that net effects as defined by Lawlor (1979) are not the *V*_*ij*_ but 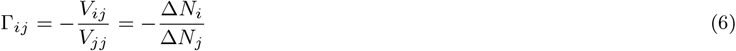 i.e. the ratio of how much species *i* responds to how much species *j* responds in the long term after a press perturbation on species *j*.

